# High-throughput Virtual Screen of Endocrine-disrupting Chemicals Identifies Disruptors of EGFR Signaling

**DOI:** 10.64898/2026.08.05.743028

**Authors:** Luke T. Jesikiewicz, Ria Marathe, Bakhtyar Sepehri, Robel Demissie, Hyun Lee, Almudena Veiga-Lopez, José A. Villegas

## Abstract

Chemical exposures during pregnancy are linked to an increased risk of pregnancy complications that contribute significantly to maternal and infant morbidity and mortality and can lead to long term health consequences for both the mother and the offspring. The placenta, a central regulator of pregnancy health, is a direct target of environmental toxicants. Epidermal growth factor receptor (EGFR), highly expressed in the placenta, regulates proliferation, migration, invasion, fusion, and cellular bioenergetics. To identify compounds of environmental concern with potential for EGFR-disrupting activity, we optimized a high-throughput virtual screening protocol for the identification of EGFR inhibitors and achieved enrichment factors of EF_1_% = 10.09, EF_5_% = 3.86, and EF_10_% = 3.0 in a benchmarking dataset. We applied this protocol to screen the Collaborative Estrogen Receptor Activity Prediction Project database, finding that top-scoring compounds were enriched for aromatic and fused-ring chemical classes, including dyes. Kinase activity assays revealed that two out of thirteen selected compounds, Vat Red 32 and Reactive Red 136, inhibited EGFR kinase activity with micromolar IC_50_ values. Additionally, pose refinement with molecular dynamics simulations characterized the binding interactions of Reactive Red 136 within the EGFR kinase domain, and functional assays in HTR-8/SVneo placental trophoblast cells showed that Reactive Red 136, but not Vat Red 32, partially attenuated EGF-mediated cell migration despite both compounds inhibiting EGFR kinase activity. Together, this study has generated an enriched dataset of candidate environmental EGFR modulators, with experimental validation confirming enrichment for EGFR-disrupting activity among the selected compounds. These results provide a valuable resource for toxicological studies.

## Introduction

The placenta is a central regulator of pregnancy health and a direct target of environmental toxicants (Gingrich *et al*., 2020). Among the receptors most relevant to placental function is the epidermal growth factor receptor (**EGFR**), which is expressed at higher levels in the placenta than in any other non-tumorigenic human tissue (Uhlén *et al*., 2015) and is required for trophoblast proliferation, differentiation, fusion, and invasion (Waye *et al*., 2025). Disruption of EGFR signaling in placental tissue has been associated with impaired trophoblast function and adverse outcomes for both mother and offspring (Waye *et al*., 2025).

A growing number of structurally unrelated environmental chemicals have been reported to interfere with EGFR signaling. For instance, bisphenol S antagonizes EGFR by binding directly to two sites within its extracellular domain (Ticiani *et al*., 2023) blocking EGF-mediated extravillous trophoblast invasion (Ticiani *et al*., 2022) and cytotrophoblast fusion (Ticiani *et al*., 2021). The organochlorine pesticide trans-nonachlor and the industrial contaminants PCB-126 and PCB-153 similarly act as competitive antagonists at the EGFR extracellular domain, displacing bound EGF (Hardesty *et al*., 2018) and dioxin-like compounds including PCB126 and TCDD have independently been shown to dock to the same extracellular subdomains (I and III), likewise preventing growth factor-induced receptor activation (Vogeley *et al*., 2022). Other environmental chemicals bearing reactive quinone or aromatic ring structures have also been reported to engage EGFR directly. The diesel exhaust component 1,2-naphthoquinone forms a covalent bond with Lys80 in the EGFR extracellular domain, activating EGFR-Akt signaling (Nakahara *et al*., 2021). To our knowledge, of the environmental chemicals characterized to date, only the herbicide atrazine has been predicted to act at the intracellular kinase domain (Hardesty *et al*., 2018), functioning as an ATP-competitive inhibitor analogous to the mechanism of clinically used EGFR tyrosine kinase inhibitors (Zubair and Bandyopadhyay, 2023). This asymmetry suggests that the kinase domain remains comparatively unexplored as a site of environmental chemical interference, despite being a druggable pocket in other contexts (Halder *et al*., 2023). Whether additional environmental chemicals act through this mechanism remains unknown.

Relatively few environmental compounds have been evaluated for EGFR-disrupting activity, given the number in commerce and human exposure. Specifically, no systematic screen has evaluated candidate chemicals specifically against the EGFR kinase domain. Existing large-scale chemical screening efforts, including Collaborative Estrogen Receptor Activity Prediction Project (**CERAPP**), have primarily relied on ligand-based approaches, such as quantitative structure-activity relationship (**QSAR**) models, that predict biological activity from molecular features alone, without addressing the target protein’s three-dimensional structure, and well suited to screening very large compound libraries efficiently (Mansouri *et al*., 2016). Structure-based docking against the EGFR kinase domain, by contrast, is a well-established strategy in oncology drug discovery, having guided the design and optimization of clinically approved EGFR tyrosine kinase inhibitors (Zubair and Bandyopadhyay, 2023), but it has not, to our knowledge, been applied to prioritize environmental chemicals for kinase-domain activity. Computational docking more broadly offers a scalable, cost-effective strategy to prioritize candidates from large chemical libraries prior to experimental testing (Hemmerich and Ecker, 2020).

In this study, we repurposed the CERAPP chemical library, originally curated for ligand-based endocrine-disruption modeling (Mansouri *et al*., 2016), to conduct a structure-based, high-throughput virtual screen of candidate compounds against the EGFR kinase domain, benchmarked against a curated set of known kinase binders and non-binders (Mysinger *et al*., 2012) Top-scoring candidates from the virtual screen were evaluated biochemically for their ability to inhibit EGFR kinase activity, and compounds confirmed as kinase inhibitors were further tested functionally in a first-trimester human trophoblast cell model to determine whether they disrupt EGF-mediated trophoblast migration and invasion, and whether these effects depend on EGFR activity. This integrated pipeline allows candidate environmental chemicals to be prioritized and validated as EGFR disruptors with direct relevance to placental cell function.

## Methods

### Datasets

Enrichment analysis was carried out using a dataset of 832 known binders and 35,442 known non-binders (decoys) of the kinase domain obtained from the DUD-E database (Mysinger *et al*., 2012). The proportion of known binders for the kinase domain of EGFR in this dataset is approximately 2.29%. For the identification of EGFR disrupting chemicals, 34,141 compounds were obtained from the Collaborative Estrogen Receptor Activity Prediction Project (CERAPP) database to screen against the kinase domains of EGFR (Mansouri *et al*., 2016).

### High-throughput Virtual Screening

LigPrep as a part of Schrödinger program (version 2023-4) was used for the preparation and minimization of compounds. Protonation states at pH 7.0 were generated via Epik, specified chiral centers were maintained, and non-specified stereoisomers and tautomers were generated. The resulting 43,904 generated structures were minimized using the OPLS_2005 forcefield.

The crystallographic structure of the kinase domain of EGFR (PDB ID: 2rgp) was taken from the Protein Data Bank. The protein structure was prepared using Protein Preparation Wizard (Schrödinger (version 2023-4). Disulfide bonds were automatically assigned, hydrogen atoms were added, protonation states were determined using PROPKA (version 3) at pH 7.0 and protein structure was minimized using OPLS_2005 force field with cutoff RMSD of 0.3 Å.

Glide in Schrödinger (version 2023-4) was used to run docking calculations of the benchmark dataset(Halgren *et al*., 2004; Friesner *et al*., 2004) with a 0.8 VdW scaling factor, a partial charge cutoff of 0.15, and flexible ligand sampling under three different setting:

1. Sample nitrogen inversions on, sample ring conformations off, enhance planarity off.
2. Sample nitrogen inversions off, sample ring conformations on, enhance planarity off.
3. Sample nitrogen inversions off, sample ring conformations on, enhance planarity on.

Epik state penalties were included in the docking score. Post-docking minimization was performed on generated poses using 5 poses per ligand. The Glide grid for this site was centered at the enzyme binding pocket at coordinates (17.5, 35.08, 91.66) with 10 Å edges specified for the inner box and 32 Å edges specified for the outer box. Virtual screening of the CERAPP database was carried out with the above settings, with sample nitrogen inversions off, sample ring conformations on, and enhance planarity off. Ligands with more than 500 atoms and 100 rotatable bonds were excluded from scoring. Receiver operator curves for each benchmarking run were generated by calculating the true positive rate (TPR = TP / (TP + FN)) and false positive rate (FPR = FP / (FP + TN)) for compounds with a docking score below a threshold value of the glide score, where TP is the number of true positives, FN is the number of the number of false negative, FP is the number of false positives, and TN is the number of true negatives. The threshold value was varied by 0.01 from and covered the range of all scores. For the screening dataset, compounds were ranked by glide score and the top 300 compounds, comprising 7.7% of the total library, were selected as the enriched dataset.

### Kinase Activity Assays

EGFR kinase activity was measured using the ADP-Glo Kinase Assay (Promega, Madison, WI, USA) according to the manufacturer’s instructions with minor modifications. Test compounds (Alfa Chemistry) were prepared as 2.59, 5, or 10 mM stock solutions in DMSO and serially diluted in 100% DMSO to generate 50× working stocks. These were further diluted into kinase reaction buffer consisting of 40 mM Tris-HCl (pH 7.5), 20 mM MgCl_2_, 2 mM MnCl_2_, 50 μM DTT, and 0.1 mg/ml BSA. Recombinant human EGFR kinase (SignalChem, Part No. E10-112G-10) was diluted in kinase reaction buffer to a 2.5× working concentration of 4 ng/μL. ATP was prepared at a 5× concentration of 250 μM in kinase reaction buffer. Poly(Glu_4_,Tyr_1_) substrate (SignalChem, Part No. P61-58) was supplied at a 5× concentration of 1 μg/μl. For each reaction, 2 μL of 2.5× EGFR solution was combined with 1 μl of 5× compound solution and incubated for 5 minutes at room temperature. The reaction was initiated by adding 2 μL of a 2.5× Poly(Glu_4_,Tyr_1_)/ATP mixture, resulting in final concentrations of 1.6 ng/μl EGFR, 0.2 μg/μl substrate, and 50 μM ATP. Compound concentrations ranged from 0.0005 to 10 μM (3-fold serial dilutions) or 0.195 to 100 μM (2-fold serial dilutions). All reactions were performed in triplicate in white 384-well microplates. Reactions were incubated at 25 °C for 1 hour. Following incubation, 5 μl of reaction mixture was combined with 5 μl of ADP-Glo reagent and incubated for 40 min at room temperature. Then, 10 μl of kinase detection reagent was added, followed by an additional 40-min incubation. Luminescence was measured using a POLARstar Optima microplate reader (BMG LABTECH, Ortenberg, Germany). Percent inhibition was calculated relative to control reactions without compound, and IC_50 v_alues were determined by fitting the data to a three-parameter Hill equation using SigmaPlot v15.0 (Systat Software, San Jose, CA, USA).

### Classical Molecular Dynamics Simulations

The lowest scoring pose from all three ligands with the above docking protocol was optimized using Gaussian09W (Revision A.02) (Frisch *et al*., 2009) with the B3LYP functional and a 6-31G(d) basis set. The structure of the EGFR kinase domain was taken from the PDB (access code 2RGP) (Xu *et al*., 2008). Missing residues were added to the structure of chain A using the ModLoop server (Fiser and Sali, 2003). The protonation state of charged residues were determined using the H++ webserver at pH 7 and 0.15 μmol salinity (Gordon *et al*., 2005).

Classical molecular dynamics (MD) simulations were carried out using the pmemd module of the GPU-accelerated Amber 22 package (Case *et al*., 2022). The Amber ff14SB force field (Maier *et al*., 2015) was used for standard residues, TIP3P was used for solvent water molecules and ions. Using the Merz-Singh-Kollman scheme the RESP charge fitting on electrostatic potential generated at the HF/6-31G(d) level of theory for each ligand, utilizing the generalized Amber force field (gaff) (Wang *et al*., 2004) to generate forcefield parameters for the ligands. The ligand-receptor complex was solvated in a square water box with a periodic boundary condition with the minimum distance between the receptor and the edge of the box as 12 Å. The classical MD simulations follow four stages; minimization, heating, equilibration, and production runs. We applied 30,000 steps of energy minimization with 5.0 kcal mol^-1^ Å^-2^ restraints on the receptor and ligand. Then, the system was heated to 300 K over 10,000 steps of MD at a 1fs timestep with the same 5.0 kcal mol^-1^ Å^-2^ restraints at constant volume. The system was then equilibrated over 600,000 steps of MD at a 1fs timestep, gradually releasing the harmonic restraints every 100,000 steps.

Production simulations were performed at constant pressure with the Beresden barostat and 300 K (Lagevin thermostat) with a 2 fs timestep with 0.1 kcal mol^-1^ Å^-2^ restraints on the backbone atoms of the GR only. Production runs were performed for 150 ns in triplicate, employing the SHAKE algorithm for H atoms, the Particle-Mesh Ewald method (Darden *et al*., 1993) for long-range electrostatic effects, and a 10.0 Å cutoff for electrostatic interactions. Frames were written to the file every 5000 steps (15,000 frames per simulation).

CPPTRAJ (Roe and Cheatham, 2013) was utilized for RMSD calculations as well as interaction distances and frequencies. Clustering analysis based on the root-mean-square deviation (**RMSD**) of receptor backbone was carried out using the CPPTRAJ module to identify the most populated ligand conformation in the MD simulations of all three replicas. H-bonds are specified as having a donor-acceptor atom distance of <3.0 Å and a bond angle of >140°. H-bond frequency is reported as number of frames specifying H-bond criteria across the three triplicate runs of MD simulation divided by the total number of simulation frames. 3D renderings were created using Pymol (The PyMOL Molecular Graphics System, Version 3.0 Schrödinger, LLC).

### MM(GB)SA Binding Energy Calculations

Binding energy calculations were computed using the MPI implementation of the mmgbsa.py (Miller *et al*., 2012) module in AmberTools with igb=8, surface tension set to 0.0072. Energetic sampling for MM(GB)SA was done on the last 100ns of each trajectory with 0.25 ns fragments (400 total frames). Normal mode entropy calculations were performed on 8 frames within the last 100ns of each trajectory. Reported MM(GB)SA energies, and per-residue energy decomposition energies are the average energy across all three triplicate runs.

### HTR-8/SVneo Cell Culture

HTR-8/SVneo cells, a first-trimester extravillous trophoblast-derived cell line that migrates and invades in response to EGF through EGFR activation, were used to test EGFR antagonism for the screened chemicals as previously described (Ticiani *et al*., 2023). HTR-8/SVneo were cultured in complete culture media consisting of Dulbecco’s modified Eagle’s medium/F12 medium (124000-024, Millipore Sigma) supplemented with 10% of fetal bovine serum (FBS), 2 mM L-glutamine, 10 mM HEPES, 100 IU/ml penicillin, and 1% antibiotic-antimycotic (15240062, Gibco). Cells were cultured in 5% CO_2_ at 37 °C.

### CellTiter-Glo Cell Viability Assay

To determine whether a subset of the screened compounds in the kinase assay werecytotoxic to HTR-8/SVneo cells, cell viability was assessed using the CellTiter-Glo luminescent cell viability assay kit (G8091, Promega) as previously described (Ticiani *et al*., 2023). Briefly, cells were seeded in opaque plates at 10,000 cells per well overnight, then exposed to vehicle control (0.1% DMSO), Vat Red 32 (**VR32**; ACM2379773, Alfa Chemistry), or Reactive Red 136 (**RR136**; ACM83137159, Alfa Chemistry) at concentrations of 0.0001, 0.001, 0,01, 0.1, 1, and 10 μM for 24 h. All treatments were performed at least in triplicate. Following exposure, plates were equilibrated to room temperature for 30 min, and cells were then incubated with 100 µl of CellTiter-Glo per well for 10 min. Luminescence was recorded using the BioTek Cytation 1 Cell Imaging Multimode Reader (1623200, Agilent) Results were expressed as luminescent units (LUM).

### Placental Cell Migration

A scratch wound cell migration assay was utilized to assess the effects of VR32 and RR136 on HTR-8/SVneo cell migration. Cells were seeded into a clear flat-bottom 96-well plate at 30,000 cells in 100 μl per well and allowed to adhere overnight. Exposure media containing vehicle control (0.1% DMSO), VR32, or RR136 at concentrations of 0.0001, 0.001, 0,01, 0.1, and 1 μM was prepared prior to plate scratching. Scratch wounds were created using the Incucyte 96-well woundmaker tool (4563, Sartorius). Medium was then aspirated, and cells were washed twice with complete media. After washing, 100 μl of exposure media was added to each well, and the plate was placed in the Incucyte S3 live-cell analysis system (4647, Sartorius) and imaged every 2 h for 24 h. Wound width per well was quantified at each timepoint using the Incucyte scratch wound analysis software module (9600-0012, Sartorius) and normalized to the initial width (0 h). Percent wound closure was calculated as: ((WW_t_ - WW_i_) / WW_i_)) * 100, where WW_t_ is the wound width at timepoint t, and WW_i_is the initial wound width. Data were analyzed in GraphPad. To evaluate if VR32 and RR136 affect epidermal growth factor (**EGF**)-mediated cell migration, the protocol above was modified as follows. After washing, cells were pre-treated for 15 min with 50 μl of vehicle control, 1 μM VR32, or 1 μM RR136 exposure media. Following pre-treatment, 50 μl of vehicle control, 30 ng/ml human EGF (E9644, Sigma Aldrich), 1 μM VR32, 1 μM RR136, 1 μM VR32 + 30 ng/ml EGF, or 1 μM RR136 + 30 ng/ml EGF were added to the cells prior to Incucyte live-cell imaging as described before. All treatments were performed at least in triplicate.

### Statistical Analyses

Data were analyzed using GraphPad Prism (GraphPad) and are expressed as mean ± SEM. Normality was assessed using the Shapiro-Wilk test, with transformations applied as needed. Outlier values identified by ROUT method (Q = 1%) were excluded from statistical analyses. Group comparisons were performed by Brown-Forsythe and Welch’s one-way ANOVA followed by Dunnett’s multiple comparisons test for normally distributed data, or by the Kruskal-Wallis test with Dunn’s multiple comparisons test when normality could not be achieved. A P value < 0.05 was considered statistically significant.

## Results and Discussion

EGFR is a cell-surface receptor which comprises an extracellular domain that responds to epidermal growth factor (**EGF**) and an intracellular domain which functions as a tyrosine kinase (Wee and Wang, 2017; Chen *et al*., 2016). Binding of native ligands such as EGF to the extracellular domain of EGFR induces receptor homodimerization, as well as heterodimerization with other receptor types. This, in turn, activates the receptor tyrosine kinase domain through the transphosphorylation of tyrosine residues within the kinase domain. Consequently, this process initiates downstream signaling pathways that are responsible for cell proliferation, cell cycle progression, cell division, motility, invasion, and adhesion (Mitsudomi and Yatabe, 2010; Marshall, 2006). Given that the kinase domain has well-known susceptibility to inhibition by a diverse set of chemical structures, we reasoned that chemical exposures could have the potential to disrupt EGFR signaling. We therefore set out to apply a computational screening approach to identify a chemical library of compounds with a significant likelihood of EGFR disrupting potential.

### Enrichment analysis

The Glide software is a widely used tool for computational docking with a competitive success rate for the identification of candidate binders. Glide calculations are customizable by varying several parameters, including the extent of conformation sampling. The high-throughput virtual screen (**HTVS**) performs docking calculations with lowest degree of sampling to enable the processing of large compound libraries within a reasonable time frame. Additionally, other parameters allow for setting the configuration sampling of the ligand molecules, including sampling nitrogen inversions, sampling ring conformations, and optimizing planarity of ring structures. However, it is not obvious which set of parameters is likely to give the best results for a particular target. Optimal settings are often system dependent and therefore it is advisable to conduct benchmarking calculations (Jain and Nicholls, 2008). To ensure the HTVS screening protocol in Glide that was best capable of enriching a library of candidate compounds, we carried out a benchmarking procedure using a set of known binders and non-binders to the EGFR kinase domain obtained from the Directory of Useful Decoys, Enhanced (DUD-E) using three different parameter settings (Mysinger *et al*., 2012).

As docking scores are well-known to correlate poorly with experimentally calculated values of binding affinity, no inference is made between docking scores and binding affinities. Instead, a cut-off value is selected as the threshold value and all compounds with a score below this value are labeled as predicted binders. This subset of the library constitutes an enriched dataset which is expected to contain a higher percentage of true binders than the overall dataset. Given a set of known binders and decoys, True Positive Rates (TPR) and False Positive Rates (FPR) are calculated for compounds below the threshold value and are plotted to yield a receiver operator curve, for which the area under the curve (**AUC**) provides an indication of the enrichment capability of the virtual screening procedure. An AUC value of 0.5 indicates no enrichment capability, while a value of 1.0 would indicate the model is able to distinguish perfectly between binders and decoys. For the EGFR kinase domain benchmark dataset, the greatest AUC value (0.747) was obtained when sampling ring conformations and turning off enhanced planarity (**Fig. 1**), indicating that the procedure is best able to provide enrichment capabilities of the combined dataset. Moreover, we found substantial early enrichment of true binders at low docking score threshold values, with enrichment factors of EF_1_% = 10.09, EF_5_% = 3.86, and EF_10_% = 3.0 (**Fig. 2**).

**Figure 1.**
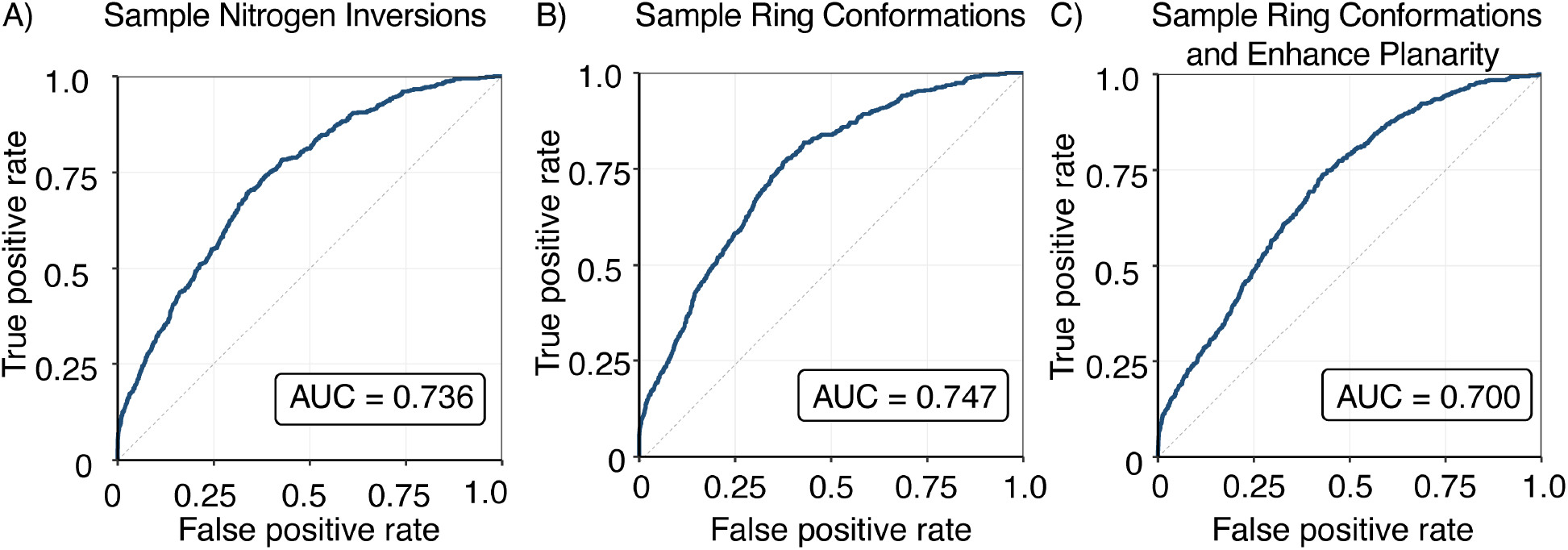
Receiver operator curves for docking results of benchmark at different parameters settings. A) Sample nitrogen inversions. B) Sample ring conformations. C) Sample ring conformations and enhance planarity.

**Figure 2.**
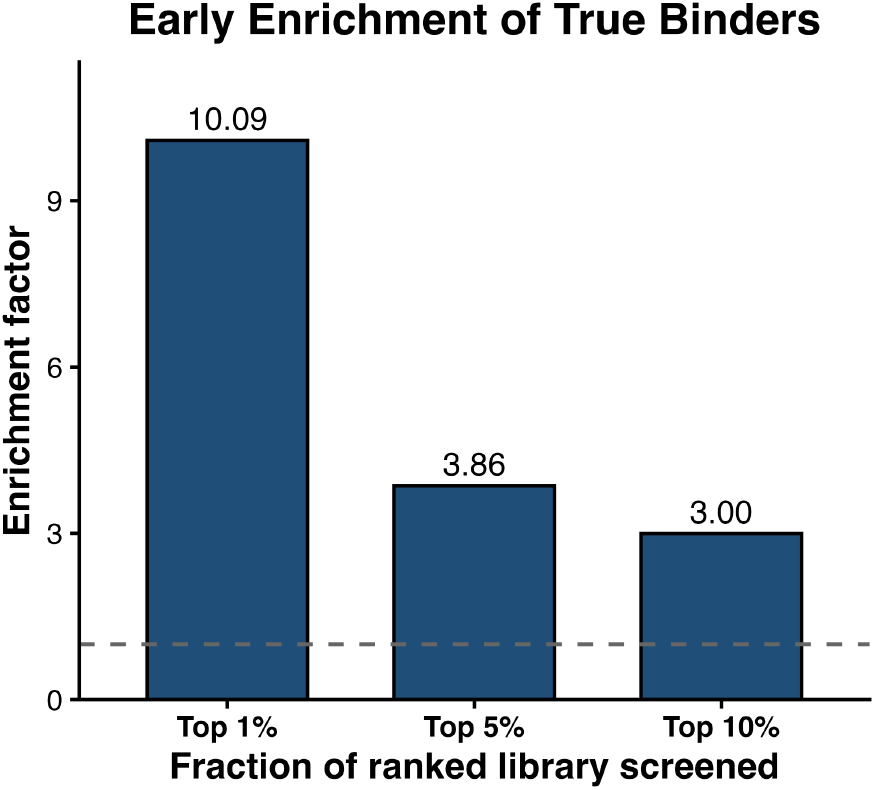
Enrichment factors of top 1%, 5%, and 10% of compounds by Glide score. Subset of top 1% scoring compounds contained 10.09x the number of true binders than the benchmark dataset, while the top 5% and 10% contained 3.86x and 3.00x respectively.

### Library Enrichment by Virtual Screening of CERAPP database

We applied the selected HTVS methodology to screen compounds in the CERAPP database for potential EGFR kinase domain inhibitory activity. The top 300 scoring compounds, constituting 9.2% of the database, ranged in scores from –10.479 to –7.749. We hypothesized that this dataset is enriched in compounds with the potential to inhibit EGFR kinase activity as this docking score produced a true positive rate of 0.20 and a false positive rate of 0.05, indicating that below this threshold docking score value the procedure is able to identify true binders at a higher rate than producing false positives. Additionally, the benchmarking dataset below this threshold value contains 6.9% true binders, compared to 1.8% in the entire dataset, indicating an enrichment factor of 3.8.

In order to further understand overall trends within top compounds selected by the docking protocol, the top 300 and bottom 300 compounds were evaluated using the ClassyFire webserver and sorted by molecular class. The top 300 compounds had contributions from 55 unique molecular classes, with benzene and derivatives ranking top with (25.7%), followed by naphthalenes (16.3%) and anthracenes (9%). The bottom 300 compounds had contributions from 72 unique classes and similarly had benzene and derivatives as the top contributor (29.3%). The contribution of naphthalenes was significantly lower than the top 300 (down 12%) and anthracenes were not found to contribute more than 5 representative structures suggesting the docking protocol favors selection of fused cyclic compounds (**Fig. 3**). Additionally, flavonoids, pyrenes, azobenzenes, indanes, and indoles were classes unique to the top 300.

**Figure 3.**
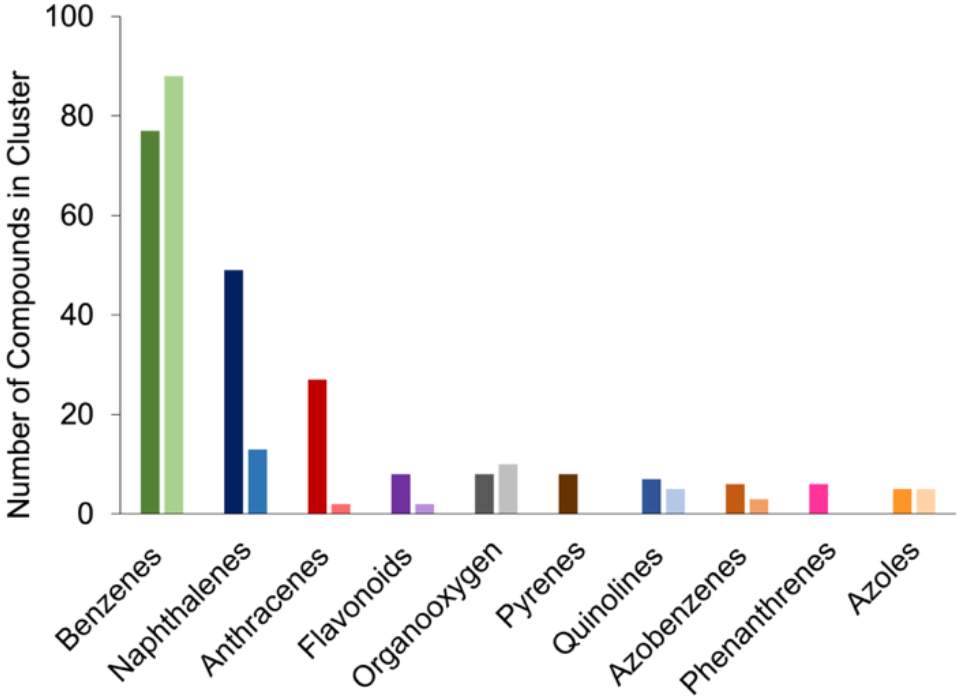
ClassyFire classification of screend compounds from CERAPP database. (Left columns) Top 300 scoring compounds. (Right columns) Bottom 300 scoring compounds.

Out of the set of 300 compounds, we selected a representative sample for experimental investigation. We chose 13 exogenous chemicals based on their structural classes and potential for occupational exposure (**Fig. 4A**). These compounds were CERAPP IDs 37430 (tautomeric form of reactive orange 5), 12519 (used in the dye production), 7854 (Disperse Yellow 42), 30321 (Chlorazol Blue RW), 30120 (Vat Red 32), 46689 (Reactive Red 136), 25137 (Novaluron), 26513 (NAPHTHOL AS-BR), molecule 24520 (Hexaflumuron), 24883 (Lufenuron), 10931 (Benzanthrone), 30239 (Solvent yellow 44), and 11276 (1,3-diphenylbenzene). CERAPP 37430 (CAS 23532-29-8) is the tautomeric form of reactive orange 5 (Jinfeng Shi *et al*., 2013) and is characteristic of anionic azo dyes used in textile industry. CERAPP ID 12519 (CAS 119-40-4) is an intermediate used in dye production. Disperse Yellow 42 is a dye that has been produced on a large scale. It is estimated that only in 1990, about 10,000 tons of this molecule was produced. Novaluron, Hexaflumuron, and Lufenuron, are insecticides. Benzanthrone is a dye intermediate for anthraquinone-based dyes, while Solvent yellow 44 is a pigment. 1,3-diphenylbenzene, also known as m-Terphenyl, is an aromatic hydrocarbon used in industrial applications.

**Figure 4.**
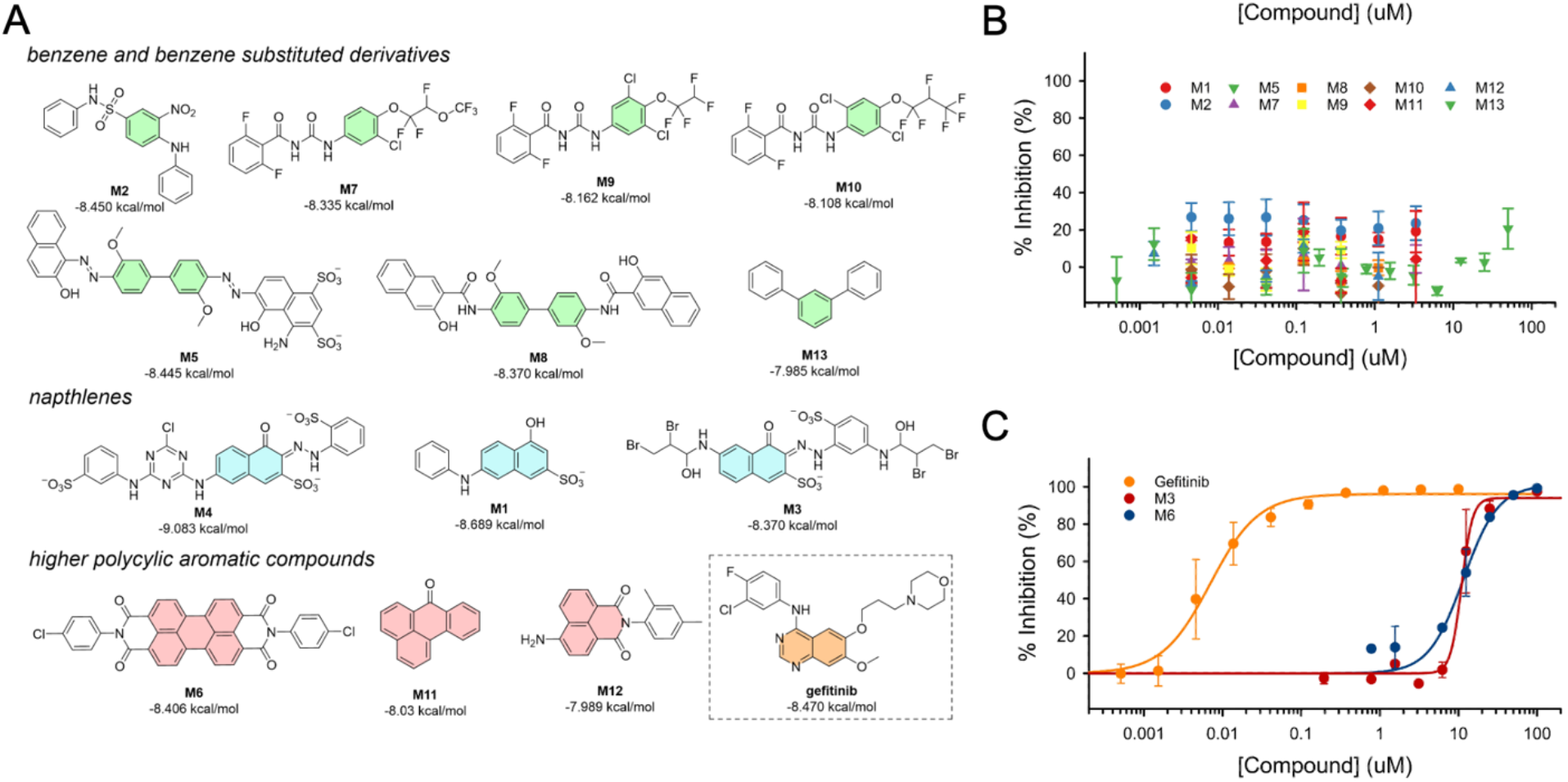
Inhibition of EGFR kinase activity by selected compounds. Dose–response inhibition curves were generated using the ADP-Glo™ kinase assay to evaluate the effects of Gefitinib (positive control), Vat Red 32, Reactive Red 136, and ten additional compounds (M1, M2, M5, M7–M13) on EGFR kinase activity. Compounds were tested over a concentration range of 0.0003–100 μM, and percent inhibition was calculated relative to vehicle-treated controls. Data points represent the mean ± SD of replicate measurements. Gefitinib exhibited potent inhibition of EGFR with an IC_50_ value of 0.0070 ± 0.0008 μM. Among the test compounds, Vat Red 32 and Reactive Red 136 demonstrated concentration-dependent inhibition with IC_50_ values of 11.0 ± 0.58 μM and 11.6 ± 2.4 μM, respectively. Compounds M1, M2, M5, M7–M13 showed minimal or no concentration-dependent inhibition within the concentration range tested and did not yield reliable IC_50_ values. Data points represent mean ± SD from two replicate measurements. Solid lines represent fits of the inhibition data to a three-parameter Hill equation used for IC_50_ determination.

### Kinase Activity Assay

Thirteen compounds selected from the virtual screening campaign were evaluated for EGFR kinase activity using the ADP-Glo kinase assay, in which the luminescence signal is proportional to kinase activity. First, initial test was performed over a concentration range of 0.0005 – 10 μM. Eleven compounds (M1, M2, M4 – M13) showed little or no inhibition across the tested concentration range, with enzyme inhibition generally remaining below 30% (**Fig. 4B**). In contrast, two compounds, Vat Red 32 (M3) and Reactive Red 136 (M6), exhibited concentration-dependent inhibition at the upper end of the tested range. For these two compounds, we repeated the kinase activity assays using a higher concentration range and observed dose-responsive behavior, with Vat Red 32 having an IC_50_ of 11.0 μM and Reactive Red 136 having an IC_50_ of 11.6 μM (**Fig. 4C**). Overall, two of the thirteen compounds exhibited micromolar EGFR inhibitory activity, indicating that the virtual screening workflow was effective in prioritizing compounds with measurable kinase inhibitory activity.

### Molecular Dynamics Simulations

To refine the poses obtained from computational docking and gain an understanding of the intermolecular interactions resulting in EGFR inhibition, we carried out molecular dynamics simulations of the active compounds (**Fig. 5**).

**Figure 5.**
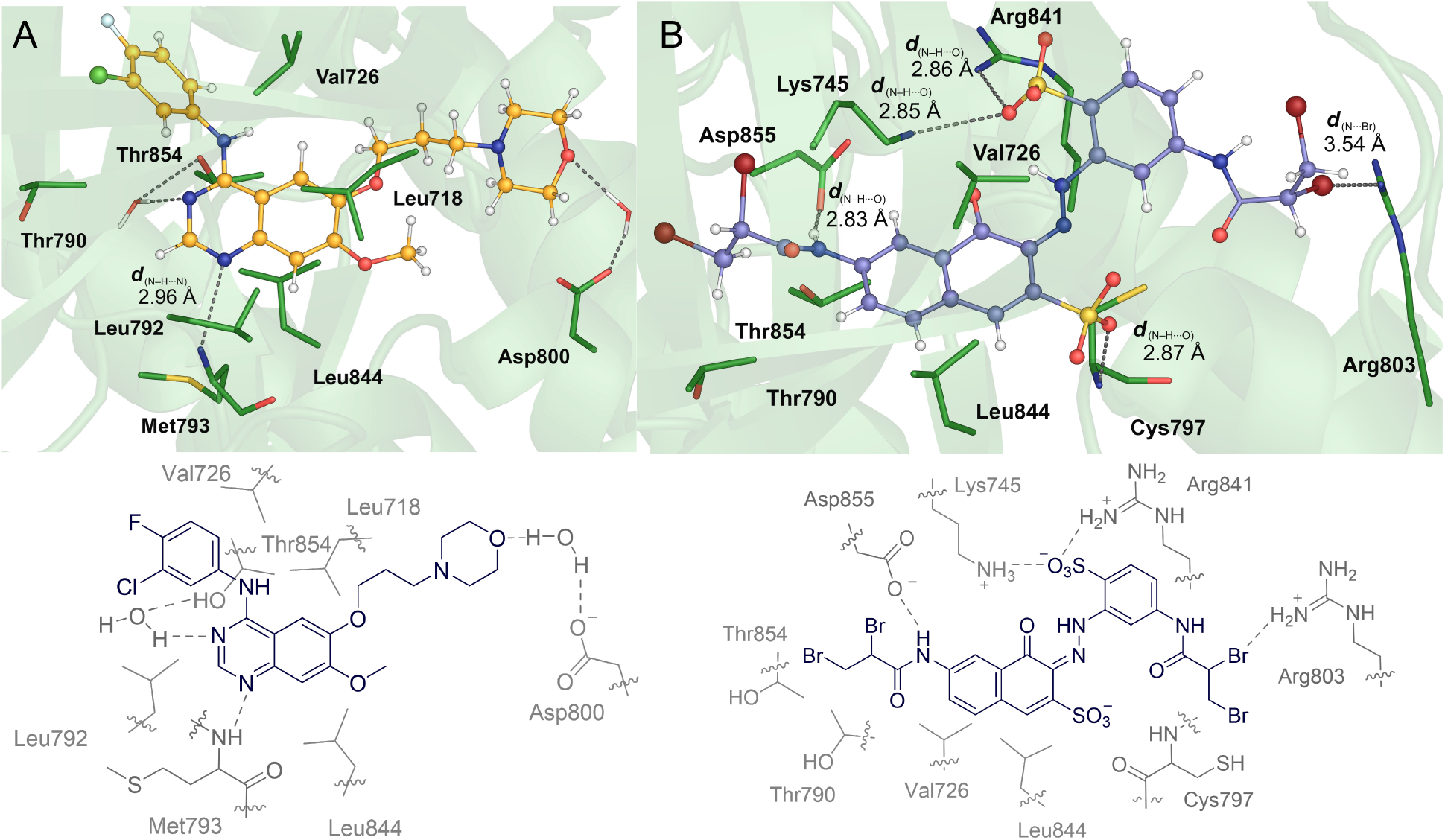
Binding pocket interactions of ligands determined from molecular dynamics simulations. (A) Gefitnib. (B) Reactive red 136 (**M6**).

Reactive red 136 (**RR136**; **M6**) was found to have a persistent electrostatic interaction between bromine and Arg803 and both bromine atoms pointed towards the exterior of the binding pocket. The secondary bromine is within 4 Å of Arg803 in 66% of frames and the primary bromine is within that same distance in 40% of frames. The most persistent hydrogen bonding interaction was between Asp855 and the napthylamide nitrogen, occurring in 77% of frames with an average distance of 2.83 Å. Similarly, there is also a hydrogen bonding interaction with the backbone nitrogen of Cys797 and the napthylsulfonate group occurring in 42% of frames with an average distance of 2.87 Å. Hydrophobic interactions with residues Val726, Thr790, Leu799, and Leu844 provide additional stabilization. **RR136** shares interactions with Leu718, Val726, and Leu844 with gefitinib, however, the core of **RR136** is shifted up in the pocket relative to gefitinib and away from the Leu792 and Met793 residues gefitinib occupies.

### Functional Validation Studies in HTR8/SVneo Extravillous Trophoblast Cells

To validate the screened chemicals on a biological level, functional *in vitro* assays were carried out using HTR-8/SVneo cells, a placental extravillous trophoblast cell line with established EGF-mediated migration and invasion (Ticiani *et al*., 2022), critical functions needed for placental attachment to the maternal decidua (Waye *et al*., 2025). To determine if Vat Red 32 (**VR32**) and RR136 are cytotoxic to EVTs, cell viability was assessed using a CellTiter-Glo assay (**Fig. 6A-B**). Both compounds were cytotoxic only at 10 μM; thus, 1 μM was used for downstream functional assays. To determine whether VR32 and RR136 interfere with EGF-mediated cell migration, HTR-8/SVneo cells were subjected to a 24-h scratch wound assay (**Fig. 6C-F**). At 16 h, EGF increased wound closure relative to control, confirming EGF-mediated migration in this assay (**Fig. 6E, F**). Neither VR32 nor RR136 alone affected wound closure relative to control (**Fig. 6E, F**). RR136 reduced EGF-mediated migration relative to EGF alone, while migration remained above control, indicating partial rather than complete antagonism (**Fig. 6E, F**), similar to other chemicals such as bisphenol S (Ticiani *et al*., 2023) VR32 had no effect on EGF-mediated migration (**Fig. 6E, F**), despite its kinase inhibitory activity in the biochemical assay. This suggests that VR32’s *in vitro* kinase activity does not translate to a functional effect on trophoblast migration under these conditions and that VR32 could be a false-positive in the chemical screen.

**Figure 6.**
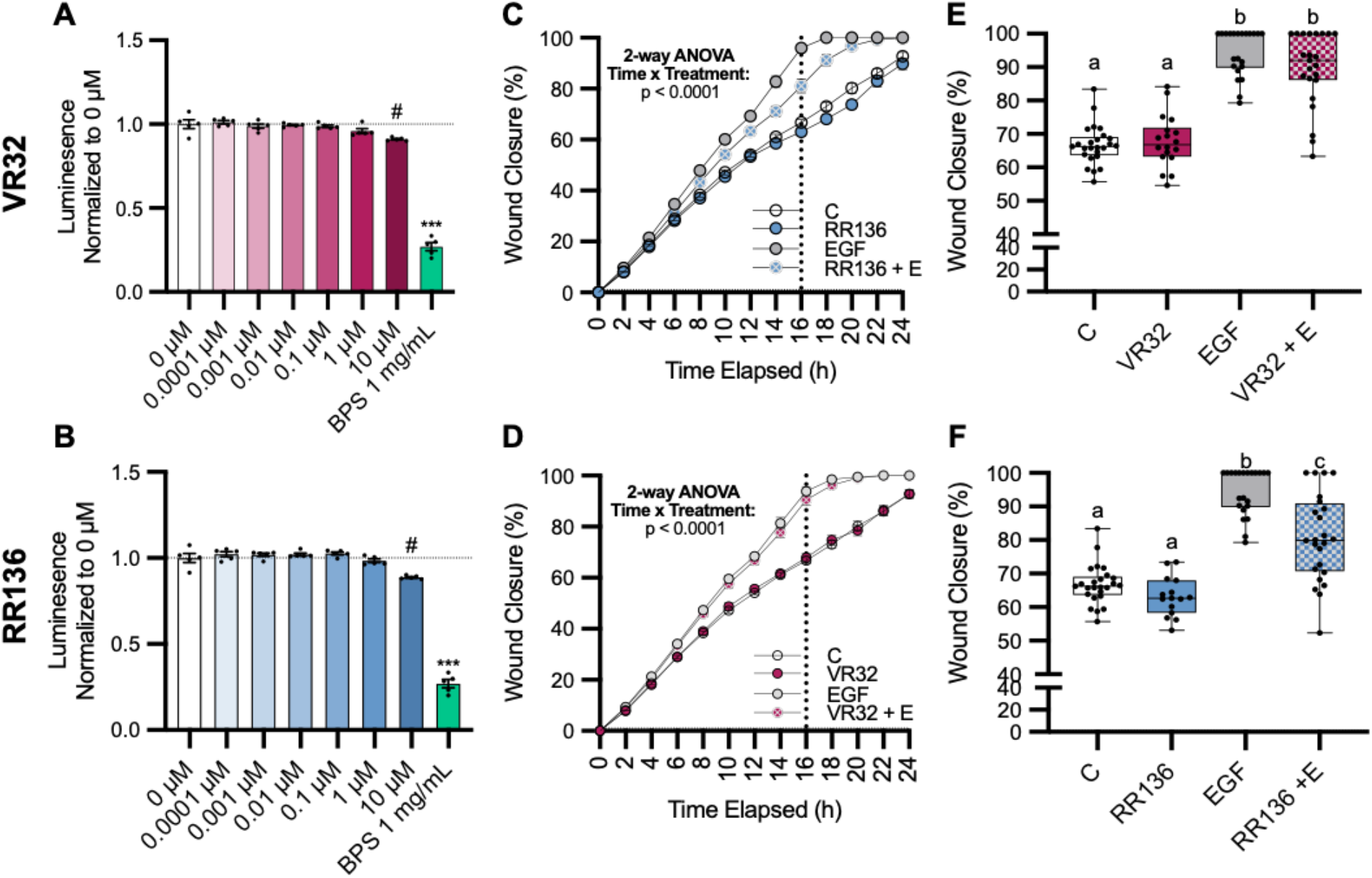
Effects of VR32 and RR136 on HTR-8/SVneo cell viability and EGF-mediated migration. (**A-B**) Cell viability was assessed via a 24h CellTiter-Glo assay in HTR-8/SVneo cells following exposure to 0, 0.0001, 0.001, 0.01, 0.1, 1, or 10 μM of VR32 (A) and RR136 (B). BPS at 1 mg/ml was used as a positive control. Brown-Forsythe and Welch’s ANOVA was used to compare groups (^#^p < 0.2, ***p < 0.0001; n = 5/dose). (**C-F**) Cell migration was assessed via a 24-h scratch wound assay following treatment with vehicle control (0.1% DMSO), 1 μM VR32, or 1 μM RR136, EGF (30 ng/ml), VR32 + EGF, or RR136 + EGF for 24 h. A 15-min pre-treatment with VR32 or RR136 was included in the EGF co-exposure groups. Wound closure over 24 h (**C-D**) and at 16 h (vertical line at 16h **C-D**; **E-F**) is shown for VR32 and RR136 (2-way ANOVA; *p < 0.0001; n = 22-25/dose). Letters differentiate groups with statistical significance (ANOVA; p < 0.001).

## Conclusion

This study establishes an integrated computational to functional pipeline, combining structure based virtual screening, biochemical kinase confirmation, and functional validation in placental cells, for identifying environmental chemicals that disrupt the EGFR kinase domain, a druggable site well characterized in oncology but comparatively unexplored for environmental chemical interference. Applying this pipeline identified Reactive Red 136 as a novel inhibitor that partially antagonizes EGF-mediated trophoblast migration, demonstrating that computational hits can be carried through to a functional, EGFR dependent outcome.

## Acknowledgments

Research reported in this publication was supported by the National Institute of Environmental Health Sciences of the National Institute of Health (R01ES035691 and P30ES027792 to A.V-L). The content is solely the responsibility of the authors and does not necessarily represent the official views of the National Institutes of Health.

## Conflict of Interest

The authors declare no conflict of interest.

## Data Availability Statement

The data that support the findings of this study are available from the corresponding author upon reasonable request.

## CRediT authorship contribution statement

**Luke T. Jesikiewicz:** Conceptualization, Methodology, Software, Formal analysis, Investigation, Data curation, Writing – Original Draft, Writing – Review & Editing, Visualization. **Ria Marathe:** Methodology, Investigation, Formal analysis, Data curation, Writing – Original Draft, Writing – Review & Editing, Visualization. **Bakhtyar Sepehri:** Methodology, Software, Formal analysis, Investigation, Data curation, Writing – Original Draft, Writing – Review & Editing. **Robel Demissie:** Investigation, Formal analysis. **Hyun Lee:** Investigation, Formal analysis. **Almudena Veiga-Lopez:** Conceptualization, Methodology, Resources, Writing – Original Draft, Writing – Review & Editing, Supervision, Project administration, Funding acquisition. **José A. Villegas:** Conceptualization, Methodology, Software, Formal analysis, Resources, Writing – Original Draft, Writing – Review & Editing, Supervision, Project administration, Funding acquisition.

## Notes

### Competing Interest Statement

The authors have declared no competing interest.

